# *Brucella* suppress innate immunity by down-regulating STING expression in macrophages

**DOI:** 10.1101/2019.12.31.891051

**Authors:** Mike Khan, Jerome S. Harms, Yiping Liu, Jens Eickhoff, Jin Wen Tan, Tony Hu, Fengwei Cai, Erika Guimaraes, Sergio C. Oliveira, Richard Dahl, Delia Gutman, Glen Barber, Gary A. Splitter, Judith A. Smith

## Abstract

Brucellosis, caused by *Brucella* bacteria species, remains the most prevalent zoonotic disease worldwide. *Brucella* establish chronic infections within host macrophages despite triggering cytosolic innate immune sensors, including Stimulator of Interferon Genes (STING), which potentially limit infection. In this study, STING was required for control of chronic *Brucella* infection *in vivo*. However, early during infection, *Brucella* down-regulated STING mRNA and protein. Down-regulation occurred post-transcriptionally, required live bacteria, the *Brucella* type IV secretion system, and was independent of host IRE1-RNase activity. Rather, *Brucella* induced a STING-targeting microRNA, miR-24-2. Furthermore, STING downregulation was inhibited by miR-24 anti-miRs and in *mirn23a* locus-deficient macrophages. Failure to suppress STING expression in *mirn23a*^−/−^ macrophages correlated with diminished *Brucella* replication, and was rescued by exogenous miR-24. Anti-miR-24 potently suppressed replication in wild type, but much less in STING^−/−^ macrophages, suggesting most of the impact of miR-24 induction on replication occurred via STING suppression. In summary, *Brucella* sabotages innate immunity by miR-24-dependent suppression of STING expression; post-STING activation “damage control” via targeted STING destruction may enable establishment of chronic infection.

**Author summary:** Cytosolic pattern recognition receptors, such as the nucleotide-activated STING molecule, play a critical role in the innate immune system by detecting the presence of intracellular invaders. *Brucella* bacterial species establish chronic infections in macrophages despite initially activating STING. STING does participate in the control of *Brucella* infection, as mice or cells lacking STING show a higher burden of *Brucella* infection. However, we have found that early following infection, *Brucella* upregulates a microRNA, miR-24, that targets the STING messenger RNA, resulting in lower STING levels. Dead bacteria or bacteria lacking a functional type IV secretion system were defective at upregulating miR-24 and STING suppression, suggesting an active bacteria-driven process. Failure to upregulate miR-24 and suppress STING greatly compromised the capacity for *Brucella* to replicate inside macrophages. Thus, although *Brucella* initially activate STING during infection, the ensuing STING downregulation serves as a “damage control” mechanism, enabling intracellular infection. Viruses have long been known to target immune sensors such as STING. Our results indicate that intracellular bacterial pathogens also directly target innate immune receptors to enhance their infectious success.

## Introduction

*Brucella* spp. are Gram-negative, facultative intracellular α-proteobacteria which cause the zoonotic disease brucellosis (1, 2). Human brucellosis is characterized by an acute undulating fever accompanied by flu-like myalgias before developing into a chronic disease, with long-term pathologies such as sacroiliitis, arthritis, liver damage, meningitis, and endocarditis (3). Brucellosis in animals often causes orchitis and sterility in males and spontaneous abortions in females, leading to profound economic loss worldwide (4). During chronic infection, *Brucella* live and replicate within macrophages and other phagocytes. This intracellular localization renders the organism refractory to even prolonged multiple antibiotic treatments, and relapses occur in 5-10% of cases (3). In the U.S., brucellosis has been largely controlled through vaccination of livestock with live attenuated strains, though outbreaks still occur (Serpa et al. 2018, Joseph et al. 2018, Sfeir 2018). Currently, no safe and effective human vaccine exists. The mechanism(s) involved in supporting the intracellular persistence of *Brucella* remain unclear.

Innate immune responses form the first line of defense against bacterial pathogens. However, *Brucella* express multiple ‘atypical’ virulence factors, which stymie innate defenses. For example, *Brucella* spp. resist complement activation and express a weakly endotoxic “smooth” lipopolysaccharide that is a poor agonist for the innate immune sensor Toll-like receptor 4 (5). Despite sequestration in membranous compartments, *Brucella* trigger cytosolic innate immune sensors including various inflammasomes and the Stimulator of Interferon Genes (STING) (6-9). STING resides in the endoplasmic reticulum membrane and upon activation by bacterial cyclic-di-nucleotides or cyclic GMP-AMP (c-GAMP), STING translocates to peri-nuclear clusters where it co-localizes with and activates TANK binding kinase I (TBK1), which in turn phosphorylates the IFN-β regulatory transcription factor IRF3 (10, 11). In addition to Type I interferon induction, STING is essential for optimal induction of NF-κB-dependent pro-inflammatory cytokines and other host defense genes, and regulates autophagy (12). Evidence from the cancer literature also suggests STING critically supports effective CD8+ T cell adaptive immune responses (13). Previously, we have shown that STING is required for Type I interferon production in response to infection with *Brucella abortus*, and that STING contributes to control of *B. abortus* infection at 72 hours *in vitro* (6, 14).

Here, we report that STING is critical for the control of chronic *Brucella* infection (3-6 weeks) *in vivo*. However, early during infection, *Brucella* down-regulate STING (T*mem173*) mRNA expression and protein. Concurrently with STING suppression, *Brucella* induce a STING-targeting microRNA miR-24. Inhibition by anti-miR-24 or genetic deficiency of miR-24-2 leads to a significant increase in STING expression as well as augmented IFN-β production in macrophages. Inability to induce miR-24 and downregulate STING compromised *Brucella* survival in macrophages. These results suggest that *Brucella* mitigates the cost of innate immune activation by targeting STING expression.

## Results

### STING is required for chronic control of *Brucella in vivo*

In a previous study, we showed that STING is required for control of *Brucella in vitro* at 72 hours (14). At an earlier time point (24h), STING (*Tmem173*)^−/−^ macrophages displayed significantly increased *Brucella* infection (**Figure 1A**). Recently, we have also shown that STING is required for control of *Brucella* infection in mice at 1 and 3 weeks (15). To confirm these results and evaluate the role of STING in longer-term chronic *Brucella* infection (16), wild type C57BL/6 and STING^−/−^ mice were infected with wild-type *S2308 Brucella abortus* for 3 and 6 weeks. Splenic colony forming units (CFU) showed an approximately two-log difference between STING^−/−^ mice and age-matched control C57BL/6 mice at 3 weeks and ∼1.5 log difference at 6 weeks (**Figure 1B**). These data indicate that STING critically participates in the control of chronic *Brucella* infection *in vivo*.

**Figure 1.**
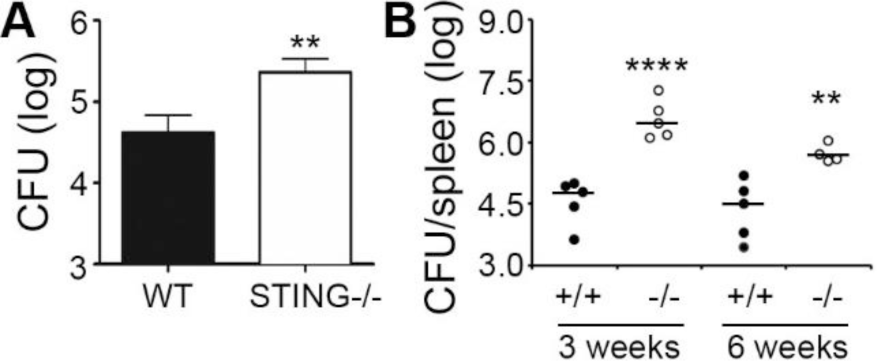
STING is required for control of acute *Brucella* infection *in vitro*, and chronic infection *in vivo*. A) Bone marrow derived macrophages from wild type C57BL/6 control (WT) or STING^−/−^ mice were infected with 10 multiplicity of infection (MOI) *B. abortus* for 24h prior to enumeration of colony forming units (CFU). Error bars denote triplicate determinations. B) Wild-type C57BL/6 (black circles, +/+) and STING^−/−^ mice (open circles, −/−) were infected for 3 or 6 weeks with 10^6^ CFU *Brucella abortus 2308* and splenocyte CFUs determined. Circles represent individual mice with 5 mice per group except the STING^−/−^ from week 6 (4 mice). Bars denote median CFU/group. Results in (A) and (B) are representative of 3 independent experiments.

### *Brucella* infection suppresses STING expression independently of IRE1 endonuclease activity and requires live bacteria and Type IV secretion

Given the requirement for STING in the control of chronic infection, it was surprising to note significant STING (*Tmem173*) mRNA down-regulation in bone marrow-derived macrophages (BMDMs) infected with wild type *16M Brucella melitensis* at 24h (published RNAseq data set in Khan et al., 2016). To confirm the RNAseq data, and determine whether other *Brucella* species down-regulate STING, *v-raf/v-myc* immortalized murine bone marrow-derived macrophages (17) were uninfected or infected with different *Brucella* species for 24 hours and STING (*Tmem173)* mRNA levels assessed via RT-qPCR (**Figure 2A**). *B. melitensis, B. abortus* and *B. suis* are human pathogens and primarily infect ruminants, cattle and swine, respectively. *B. neotomae* has been isolated from wood rats and voles but also has been isolated in human neurobrucellosis (18). The four species of *Brucella* significantly down-regulated STING mRNA compared to uninfected macrophages. STING protein levels also decreased in cells infected for 24 hours with *Brucella* (**Figure 2B**). In macrophages infected with *B. melitensis, Tmem173* mRNA down-regulation was evident by 4h post-infection (**Figure 2C**).

**Figure 2.**
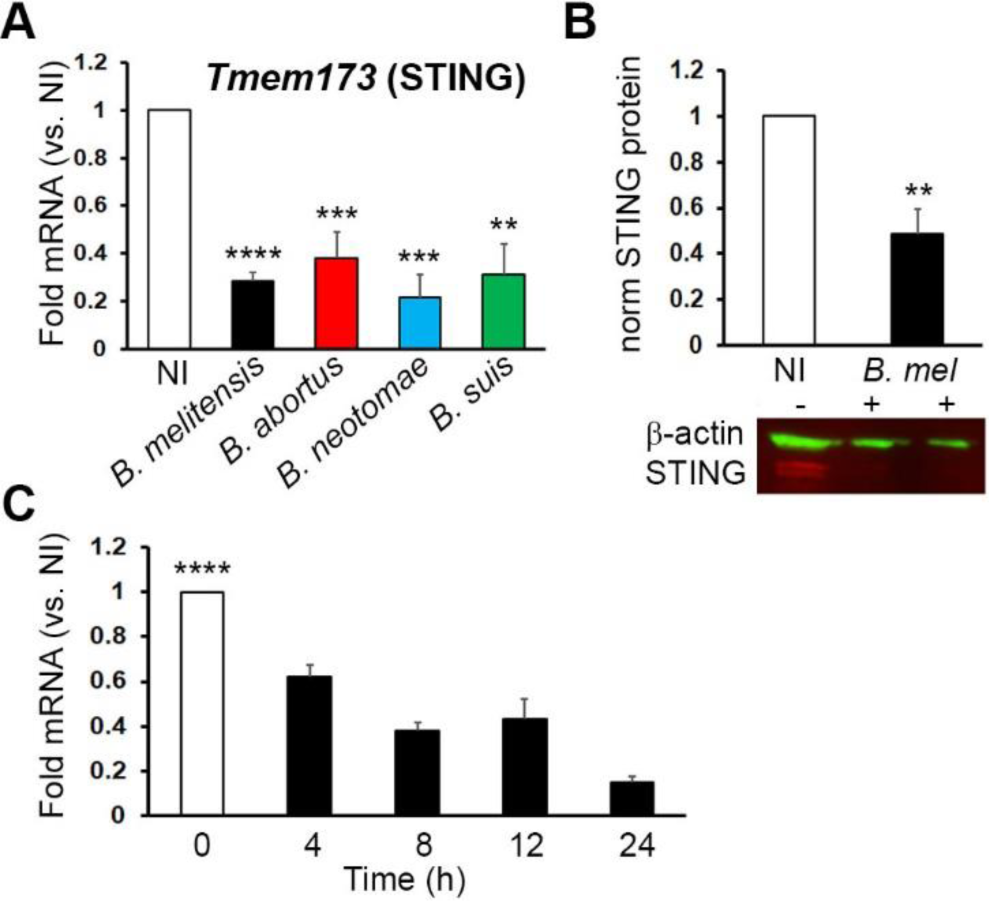
*Brucella* suppresses STING expression. A) Immortalized murine bone marrow derived macrophages were not infected (NI) or infected with *B. melitensis* 16M (black bars), *B. abortus* (red), *B. neotomae* (blue) or *B. suis* (green) as indicated at 100 MOI for 24h. Cells were lysed, RNA isolated and reverse transcribed, and relative *Tmem173* (STING) mRNA levels determined by quantitative PCR (qPCR) with normalization to 18S rRNA and uninfected controls (NI, set = 1). Results are from 25, 7, 5, and 5 independent experiments respectively, with error bars denoting SEM. P-values are vs. NI control. B) Protein expression of STING: cells were infected with 100 MOI of *B. melitensis* for 24h, and lysates resolved using SDS PAGE. STING and β-actin proteins were detected by western blot. Band fluorescence was quantitated and results are means +/- SEM of 5 independent experiments. An example western blot is shown. C) Time course: Cells were infected with 100 MOI *B. melitensis* for the times indicated and processed for RNA quantitation as in (A). Results are means +/- SEM of 6 independent experiments and for all times tested, p<0.001 vs. NI.

*Tmem173* down-regulation required live bacteria, consistent with an active bacteria-driven process (**Figure 3A**). *Brucella* with mutations in the type IV secretion system (deletion of the critical VirB2 subunit (19)) displayed an intermediate phenotype with only modest downregulation of *Tmem173*, suggesting an intact type IV secretion system (T4SS) is required for full STING suppression (**Figure 3B**). Regarding mechanism of suppression, one straightforward possibility was that *Brucella* infection suppresses the activity of transcription factors required for *Tmem173* promoter activity. To address this idea, we utilized a murine STING-promoter driven luciferase reporter (**Figure 3C**). *Brucella* infection reliably increased the activity of the STING promoter-driven construct, to a comparable extent as the heat-killed *Brucella* treatment. This result suggested *Brucella* suppressed *Tmem173* expression downstream of promoter activation.

**Figure 3.**
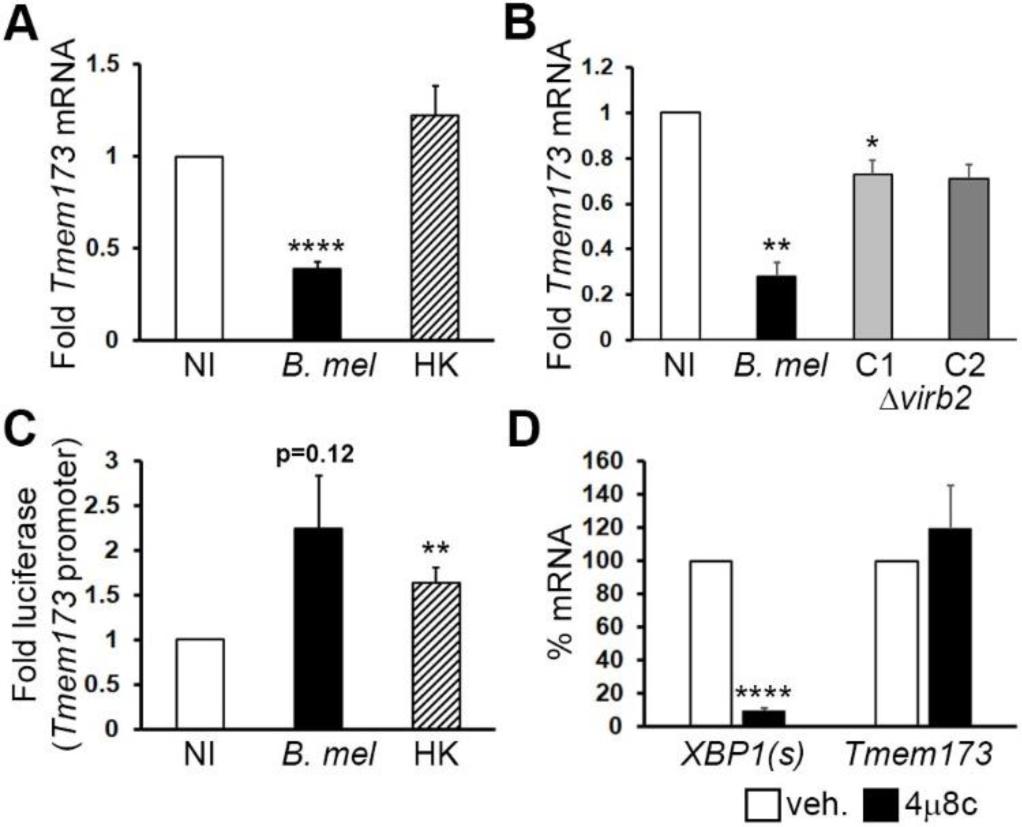
*Brucella* down-regulation of STING requires live bacteria and Type IV secretion, and is RIDD-independent. Mouse macrophages were not infected (NI) or infected with 100 MOI *B. melitensis* 16M (*B. mel*), heat-killed *B. melitensis* (HK) in (A), or 2 clones of the Δ*virb2* mutant *B. melitensis* (C1 and C2) in (B) for 24 hours prior to harvesting for RNA processing. Relative *Tmem173* mRNA levels were determined by qPCR with normalization to 18S rRNA and uninfected controls (NI set= 1) p-values are vs. NI. Results are from 21 experiments for HK and 3 independent experiments for the VirB2 mutants. C) Macrophages were transfected with a murine *Tmem173* promoter luciferase reporter. Cells were then infected with 100 MOI live (*B. mel*) or heat-killed *B. melitensis* (HK). Lysates were analyzed by dual luciferase assay. Results are from 4 and 7 experiments respectively. D) Macrophages were not treated or pre-treated with the IRE1 endonuclease inhibitor 4μ8c one hour prior to infection with *B. melitensis* (N=3 experiments). Levels of spliced XBP1 (XBP1(s)) or *Tmem173* mRNA were determined by qPCR. Untreated mRNA expression was set = 100%.

Our group and others have previously shown that *Brucella* infection induces the Unfolded Protein Response (UPR) in macrophages (20, 21). An important effector of the UPR is the transmembrane protein Inositol-requiring enzyme 1 (IRE1), which functions as both a kinase and an endonuclease. Both *B. abortus* and *B. melitenis* infections activate the IRE1 pathway (20, 22). Following activation and oligomerization, the IRE1 endonuclease cleaves 26bp from the XBP1 transcription factor mRNA, thus removing a premature stop codon in the “spliced” product (23). With prolonged endoplasmic reticulum (ER) stress, the IRE1 endonuclease changes activity to a process termed RIDD (Regulated IRE1 Dependent Decay), whereby it non-specifically degrades ER-proximal mRNAs in the secretory pathways, thus decreasing ER client load (24). To determine if RIDD degrades *Tmem173* mRNA, macrophages were pre-treated with the IRE1 endonuclease inhibitor 4μ8c (25) before infection with *B. melitensis*. As a positive control for 4μ8c efficacy, we assessed inhibition of XBP1 splicing during *B. melitensis* infection via RT-qPCR (**Figure 3D**). *Tmem173* levels were unaffected by 4μ8c pre-treatment, indicating that STING mRNA down-regulation does not occur via IRE1-dependent endonuclease activity.

### *Brucella* infection upregulates miR-24, a STING-targeting microRNA

Another hypothesis for the reduction in STING mRNA is that its mRNA is a target of microRNA (miRNA). miRNA are endogenous, small non-coding RNAs 18-25 nucleotides in length that post-transcriptionally regulate gene expression via translational inhibition and mRNA destruction (26). To search for possible miRNAs that target STING, we used the online tool TargetScanMouse to identify possible candidates. A top hit was a conserved micro-RNA miR-24, which has been shown to post-transcriptionally regulate endogenous STING in *Rattus norvegicus* epithelium cells (27). MiR-24-2 (encoded by the *mirn23a* locus) was also increased in our RNAseq data set from *Brucella*-infected macrophages (14). To confirm the effect of infection on miR-24 levels, macrophages were infected for 24 hours with *B. melitensis* (**Figure 4A**). Infected macrophages significantly and reliably increased miR-24 levels compared to uninfected cells, although the degree of induction was variable (50% up to 8-fold). To confirm the biologic relevance of this miR-24 increase, we examined expression of another predicted mRNA target BCL2-like 11 (Bim), an apoptosis facilitator (28). *Bcl2l11* mRNA levels were also significantly decreased in *B. melitensis* infected macrophages compared to uninfected cells (**Figure 4B**). *Tmem173* levels decreased over time as miR-24 increased (**Figure 4C**). Fold induction of miR-24 significantly correlated with the extent of *Tmem173* suppression (**Figure 4D**). Just as heat-killed *Brucella* failed to suppress STING, the killed *Brucella* did not induce miR-24 expression (**Figure 4E**). *Brucella* VirB2 mutants also displayed a marked defect in miR-24 induction, consistent with the failure to fully suppress *Tmem173* expression (**Figure 4F**).

**Figure 4.**
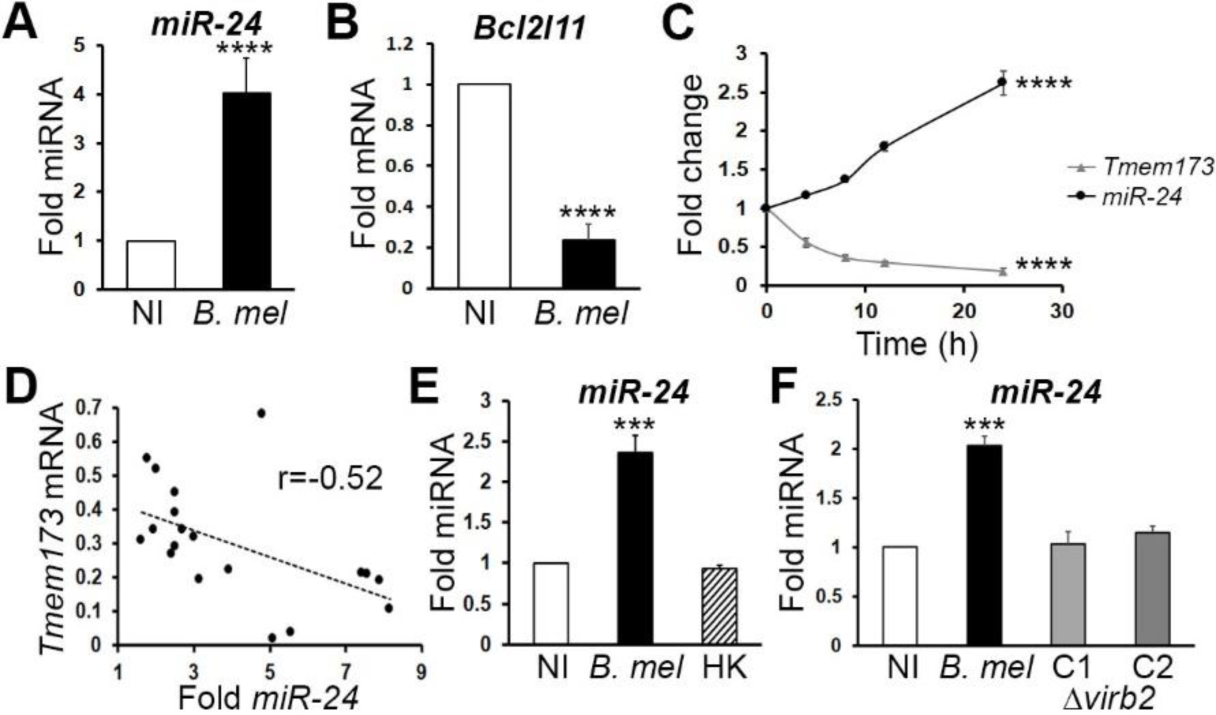
*Brucella* induces a STING-targeting microRNA miR-24. Macrophages were not infected (NI) or infected with 100 MOI *Brucella melitensis* (*B. mel*) for 24 hours before harvesting for RNA processing. mRNA or micro RNA levels were determined by qPCR with normalization to 18S rRNA or RNU6 respectively, and uninfected controls (NI set=1). A) miR-24 expression results are from 17 experiments, with error bars denoting SEM. B) *Bcl2l11* (Bim) expression is from 8 experiments. C) Time course of quantitative expression of both miR24-3p and STING (*Tmem173*) mRNA. P<0.001 for changes over time D) Correlation is from 19 experiments performed and evaluated as in (A). p=0.027 E) Comparison of live and heat killed *B. melitensis* (*B. mel* vs. HK) is from N=9. p<0.005 for *B. mel* vs. NI and HK. HK. F) Macrophages were infected with wild type *B. melitensis* or VirB2 deletion mutant clones C1 and C2 and analyzed as in (A). Results are from 3 experiments. P<0.005 for *B. mel* vs. NI and vs. ΔVirb2 clones.

The requirement for live bacteria and the type IV secretion system to induce miR-24 and suppress STING expression suggested an active, bacterially driven process, rather than a passive host response to PAMPs. Intriguingly, Ma et al. had reported that LPS suppressed STING expression via a MyD88-dependent pathway (29). The mechanism was, and remains unknown. MyD88, a critical signaling intermediary downstream of multiple Toll-like receptors, is critical for control of *Brucella* infection *in vivo* (30, 31). To determine if MyD88 contributed to *Tmem173* downregulation in *Brucella*-infected macrophages, we compared *Tmem173* and miR-24 expression in *MyD88*^−/−^ and wild type macrophages (**Figure 5A, B**). MyD88 was not required for *Tmem173* mRNA suppression or for miR-24 induction, although both were less robust in *MyD88*^−/−^ cells. Further, miR-24 induction in STING^−/−^ macrophages is similar to wild type cells, indicating mir-24 induction does not require STING expression (**Figure 5C).**

**Figure 5.**
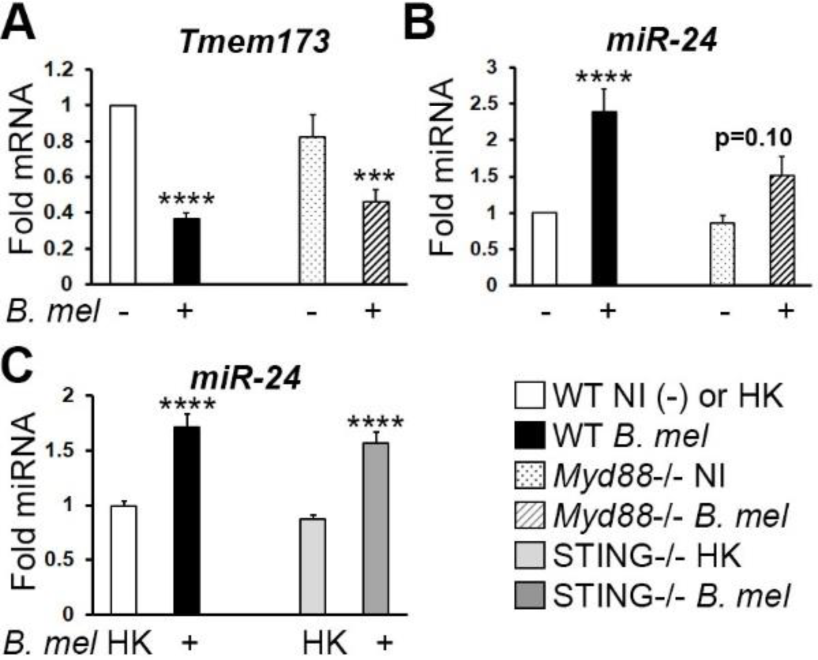
MiR-24 induction occurs in MyD88^−/−^ cells, and does not require STING. A,B) Wild type (WT) or *Myd88*^−/−^ macrophages were infected with 100 MOI *B. melitensis* (*B. mel*) for 24h and then RNA levels assessed by qPCR as above. A) *Tmem173* expression is from N=20, with normalization to WT uninfected controls within each experiment (WT NI=1). B) miR-24 expression is from N=10, normalized as in (A). C) WT or STING (*Tmem173*)−/− macrophages were infected with heat killed (HK) or live *B. melitensis* for 24h and analyzed for miR-24 expression by qPCR, with normalization to uninfected controls (NI=1), N=8. White bars: uninfected or HK-infected wild type; black bars: infected wild type; dotted bars: uninfected *Myd88*^−/−^; striped bars: infected *Myd88*^−/−^; light gray: HK infected STING−/−; dark gray: live *Brucella* infected STING^−/−^.

To confirm that miR-24 is required for the down-regulation of STING and Bim, we utilized anti-miR-24 miRNA inhibitors (**Supplemental Figure S1**). The restoration of *Tmem173* and *Bcl2l11* expression with anti-miR-24 treatment (**Figure 6A**) was consistent with the idea that miR-24 contributes to the down-regulation of these mRNAs during *Brucella* infection. STING is required for optimal *Brucella*-dependent IFN-β production in macrophages (6, 15). To determine if failure to suppress STING correlated with increased STING activity, we assessed the impact of the anti-miR-24 on IFN-β production. As shown in **Figure 6B**, IFN-β was significantly up-regulated in macrophages transfected with the miR-24 inhibitor compared to mock transfected control cells, consistent with increased STING activity. Together, these data suggest *Brucella* infection induces miR-24 to down-regulate STING.

**Figure 6.**
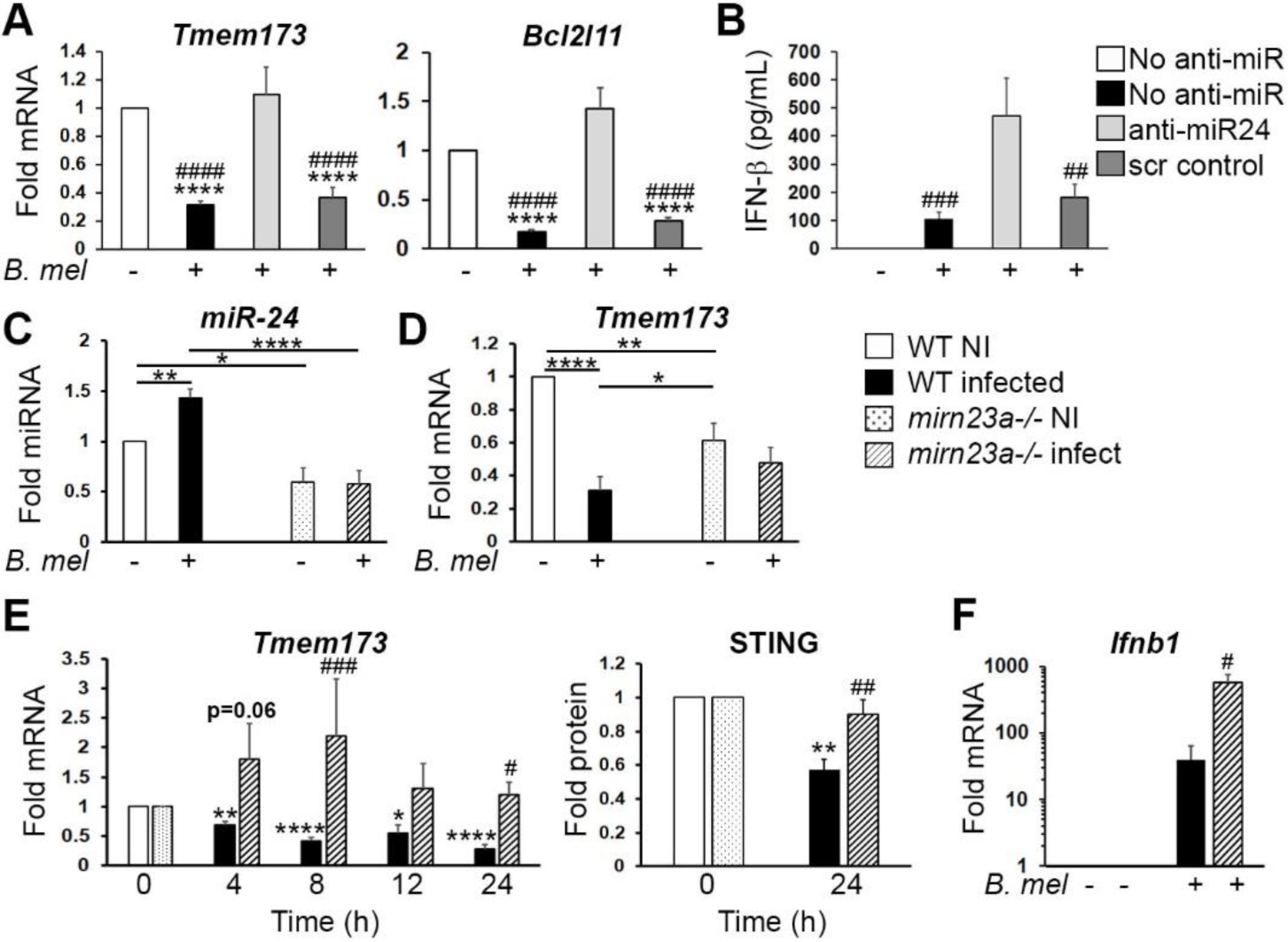
STING suppression requires miR-24 induction. A) Macrophage cells were transfected with an anti-miR24 inhibitor or control anti-miR, then infected with 100 MOI *B. melitensis* (*B. mel*) for 24h. Relative gene expression of *Tmem173* (left) and *Bcl2l11* (right) were determined via qPCR with normalization to 18S rRNA and non-infected control (NI). N=5 and 3 experiments respectively. P-values are vs. NI (*) or vs. anti-miR24 (#). (A and B) White bars are NI; black *B. mel*; light gray anti-miR-24 + *B. mel*; and dark gray scrambled anti-miR control + *B. mel*. B) IFN-β production in culture supernatant after 24h of infection was determined via ELISA. Data are from 4 experiments. C) And (D) Wild type (WT) or *mirn23a*^−/−^ macrophage cells were infected with 100 MOI *B. melitensis* for 24h and expression of miR-24 (C) *Tmem173* mRNA (D) determined as above. Results were normalized to uninfected wild type (NI=1) within each experiment. Data are from 5 and 8 experiments respectively. E) WT or *mirn23a*^−/−^ macrophages were infected with *B. melitensis* for the times indicated prior to lysis for RNA extraction. *Tmem173* levels were determined using qPCR with each genotype normalized to its own NI values (set=1). 24h data is from 9 experiments with the other time points assessed in 5 experiments. P-values for *B. mel* vs. NI in wild type cells (*) and for WT vs *mirn23a*^−/−^ infected cells (#). STING protein 24h following infection was detected using western blot with normalization to β-actin and genotype-respective uninfected controls. N=3, p-value for WT vs *mirn23a*^−/−^ infected cells (#). F) Ifnb-1 mRNA expression at 24h, normalized to uninfected control cells for each genotype. N=3. For C-F, dotted bars are uninfected *mirn23a*^−/−^ and striped bars are infected *mirn23a*^−/−^ cells.

To further evaluate the requirement for miR-24, we utilized a genetic model of miR-24 deficiency. MiR-24-3p is 100% homologous between mouse, rats and human and is expressed from two genetic loci: *mirn23a* encodes miR-23a, miR-24-2 and miR-27a and *mirn23b* encodes miR-23b, miR-24-1 and miR-27b. Our previous RNAseq data suggested bone marrow macrophages induced miR-24-2 but not miR-24-1 (Khan et al. 2016). *Mirn23a* is the predominant source of these miR-24 in blood (Kurkewich et al., 2018). *Mirn23a*^−/−^ macrophages showed decreased levels of miR-24 compared to wild type prior to infection and were deficient at miR-24 upregulation in response to *Brucella* infection (**Figure 6C**). As noted above, heat-killed *Brucella* did not induce miR-24 in either genotype. *Mirn23a*^−/−^ macrophages were unable to reliably suppress *Tmem173* expression at 24h in relation to their uninfected state, although overall levels of *Tmem173 mRNA* were decreased compared to uninfected wild type macrophages, suggesting a balance between static miR-24 and *Tmem173* levels (**Figure 6D, E).** The defect in *Tmem173* suppression in the *mirn23a*^−/−^ macrophages, more evident over time (**Figure 6E**), correlated with greatly increased *Ifnb1* induction by 24h post-infection, consistent with increased STING activity (**Figure 6F**).

### Decreased *Brucella* replication in *miR23* locus−/− macrophages

Although the data in **Figure 1** suggested that STING regulates *Brucella* infection, the biologic consequences of miR-24 induction and failure to suppress STING expression were not clear. To determine the role of the *mirn23a* locus in infection, we compared replication (CFU) in wild type vs. *mirn23a*^*−/−*^ macrophages (**Figure 7A**). Initial uptake of *Brucella* was similar between genotypes, but diverged by 8h, with lower *Brucella* CFU recovered in the *mirn23a*^*−/−*^ macrophages. This divergence maintained or increased over the course of infection through 48-72 hours. These results were consistent with a role for the miRs encoded by this locus in supporting intracellular infection. To confirm the specificity for miR-24, anti-miR24 and miR-24 mimics (**Figure 7B**) were introduced. Anti-miR24 greatly decreased the capacity of wild type but not *mirn23a*^−/−^ macrophages to control intracellular *B. melitensis* replication. In the converse experiment, addition of miR-24 mimics significantly enhanced *B. melitensis* replication in the *mirn23a*^−/−^ but not always in wild type macrophages. These data were consistent with the hypothesis that miR-24 is responsible for the decreased replication in *mirn23a*^−/−^ cells. Finally, to determine what proportion of the miR-24 effect was due to STING (vs. other miR-24 targets), STING^−/−^ macrophages were transfected with anti-miR24 or mimics prior to infection. Whereas anti-miR24 suppressed *Brucella* replication in wild type macrophages, neither mimics nor anti-miRs exerted a significant magnitude of effect on replication in STING^−/−^ cells (**Figure 7C**, anti-miRs: 5-40-fold in wild type vs 24-43% CFU difference in STING^−/−^). These epistasis results suggested STING accounts for the majority of the miR-24 effect on replication during infection. Together, these data are consistent with the hypothesis that *Brucella* induction of miR-24 suppresses STING expression to increase infectious success.

**Figure 7.**
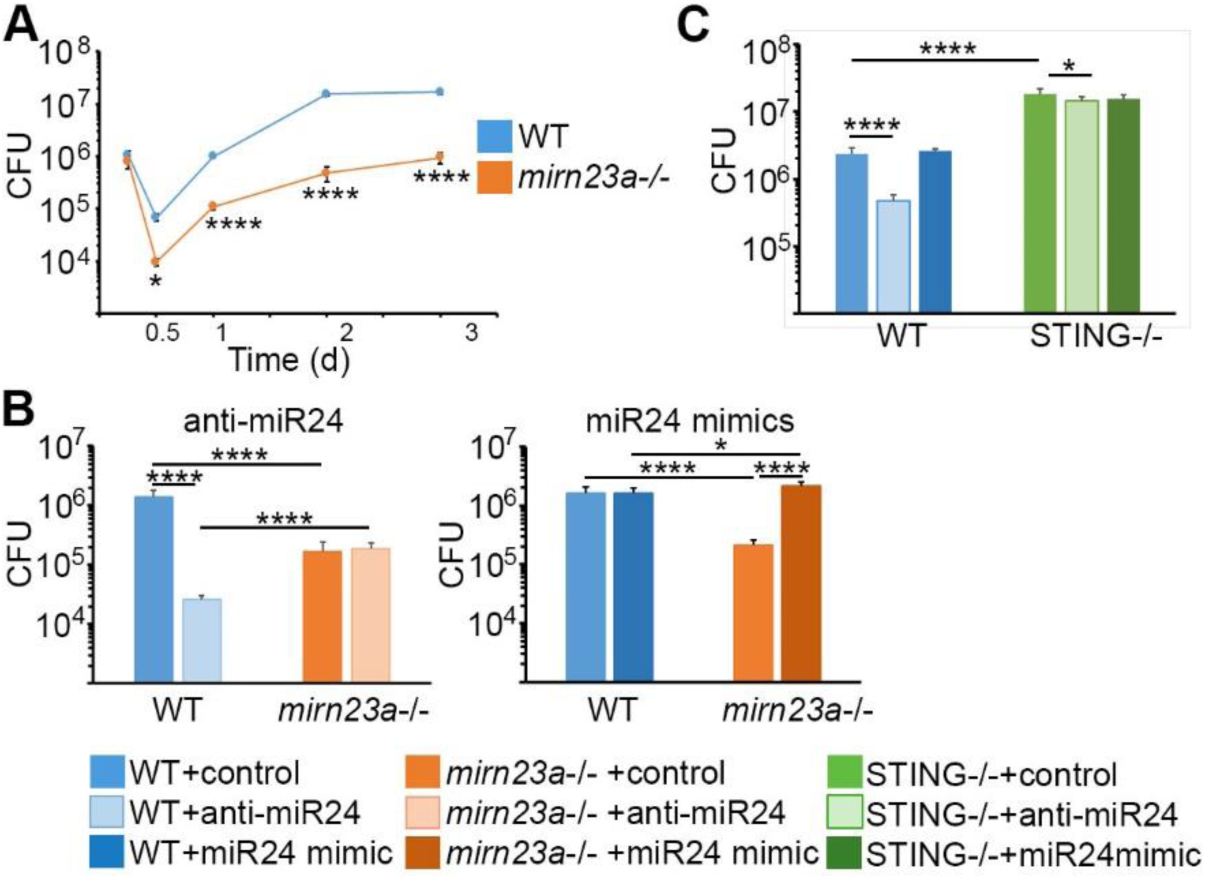
Failure to induce miR-24 inhibits *Brucella* replication. A) Wild type (WT, blue symbols) or *mirn23a*−/− macrophages (orange symbols) were infected with 100 MOI *B. melitensis* for the times indicated, lysed, and then CFU were enumerated. Error bars are standard deviations of 8 replicates and results are representative of 3 independent experiments. B) WT or *mirn23a*^−/−^ macrophages were transfected with anti-miR24 (left graph) or miR-24-3p mimic (right graph) or miR control then infected with *B. melitensis* for 24h. Cells were then lysed and CFU enumerated. Error bars are standard deviations of 8 replicates and representative of 4-5 experiments for the anti-miRs and miR-24-3p mimics respectively. Paler bars represent transfection of anti-miR24, whereas darker bars represent addition of the mimic. C) WT or STING^−/−^ macrophages were transfected with anti-miR24 or miR-24-3p mimics and then infected with 100 MOI *B. melitensis* for 24h prior to enumeration of CFU. Results are representative of 4 experiments.

## Discussion

The cytosolic DNA sensor STING plays a key role in innate immune defense via transcription of host defense genes including Type I interferons, induction of NF-κB-dependent responses and autophagy (12). *B. abortus* DNA and cyclic-di-GMP activate STING, triggering IFN-β production (6, 15). In this report, at 3 and 6 weeks post infection, STING^−/−^ mice had an almost 2-log higher burden of *Brucella* compared to age-matched wild-type counterparts, indicating that STING is ultimately required for control of chronic *Brucella* infection. Although STING protects against *Brucella*, the striking suppression of STING mRNA expression early following infection suggests *Brucella* actively sabotages this innate immune sensor to gain a foothold inside macrophages. STING suppression occurred post-transcriptionally, independently of UPR-mediated RNA decay, via upregulation of the microRNA miR-24. MiR-24 induction and STING suppression required live bacteria, and full suppression required the VirB-encoded Type IV secretion system, suggesting an active bacterial driven process rather than a simple host response to *Brucella* PAMPs. The independence of miR-24 upregulation from STING signaling supports this model.MiR-24 upregulation and STING downregulation were slightly less robust in the MyD88−/− macrophages, consistent with a minor role for MyD88 signaling. The type IV secretion system may contribute to miR-24 upregulation through secretion of a specific *Brucella* substrate or by enabling appropriate intracellular trafficking. The early divergence (4h) of STING expression between *mirn23a*^−/−^ and wild type macrophages suggests that the effect of miR-24 precedes intracellular *Brucella* replication.

Induction of the *mirn23a* gene locus during infection comes at potential cost for *Brucella* infection. In NK cells, deletion of this locus (also known as *Mirc11*) resulted in decreased ability to contain *Listeria* infection, related to diminished IFN-γ and pro-inflammatory cytokine production (32). Both IFN-γ and TNF-α have long been known to be critical for control of *Brucella* infection. However, effects of miR-24 on cytokine production may be cell-type specific. In CD4+ T cells, miR-24 was reported to target IFN-γ mRNA (33). Over-expression of miR-24 in a *Staphylococcus aureus* infection model decreased “M1” inflammatory mediator production in macrophages and enhanced “M2” marker expression, which would benefit *Brucella* (34, 35). An earlier study had also suggested modulation of macrophage polarization towards an “alternative” M2 phenotype (36). Manipulation of miR-24 levels had minimal effects on *Brucella* replication in STING^−/−^ cells, suggesting that STING is the dominant or primary target of miR-24 induction during *Brucella* infection of macrophages that impacts intracellular replication.

Recently, the ability of another chronic intracellular pathogen, *Mycobacterium tuberculosis*, to manipulate host innate responses, autophagy, and apoptosis via host miRNA has garnered much interest (37-40). In contrast, there is much less information regarding miRNA in the context of *Brucella* infection (41-43). Budak et al. investigated the miRNA expression patterns in CD4+ and CD8+ T cells from patients, and reported discrete changes with acute vs. chronic brucellosis (44, 45). Another study reported the up-regulation of miR-1981 in RAW264.7 infected macrophages and showed the interaction of that microRNA with the 3’-UTR of Bcl-2, an apoptosis regulator (46). Recently, Corsetti et al revealed several miRNA-dependent mechanisms of immune manipulation during *B. abortus* infection: upregulation of mmu-miR-181a-5p suppressed TNF-α and miR-21a-5p downregulation decreased IL10 and elevated GBP5 (43). Here, we show that miR-24 is significantly induced during infection with *B. melitensis.* Additionally, another predicted target of miR-24, Bim, a key apoptosis-regulator induced by PERK signaling and C/EBP homologous protein (CHOP) transcriptional activity, was significantly down-regulated during infection with *B. melitensis*. MiR-24 inhibition resulted in a significant recovery in both Bim and STING, indicating that miR-24 is targeting both these mRNAs during *B. melitensis* infection.

Throughout these experiments, fold-induction of miR-24 strongly correlated with STING mRNA suppression. Our previous RNAseq data set revealed upregulation of miR-24-2, encoded at the *mirn23a* locus, but not miR-24-1 from the *mirn23b* locus. Furthermore, although *mirn23a*^−/−^ cells expressed some miR-24, they were unable to upregulate its expression in response to infection. These results suggest that upsetting the balance between miR-24 and *Tmem173* levels is the critical component. The strong effects of miR-24 manipulation through mimics and anti-miRs, as well as the defects in replication in the *mirn23a*^−/−^ macrophages together support the idea that upregulation of miR-24 is important for replication early during infection. The greater replication in the *mirn23a*^−/−^ cells vs the anti-miR24 treated wild type macrophages may reflect contributions from miR-23a and miR-27a encoded by that locus (**Figure 7**).

In addition to the return of STING mRNA, miR-24 inhibition or genetic deficiency resulted in a significantly increased IFN-β response compared to uninhibited macrophages. Although initially identified in its role in viral protection, Type I interferons have recently become a topic of interest in response to many bacterial pathogens (47). During infections, the effect of Type I interferons can be protective or detrimental depending on the bacterial species. For example, Type I interferon protects mice against *Salmonella typhimurium* infection whereas Interferon-alpha/beta receptor (IFNAR)-mediated Type I interferon responses to *Francisella tularensis* and *Listeria monocytogenes* is harmful to the host (48-50). The role of Type I interferon in response to *Brucella* is currently unclear; a study in 2007 showed no difference in splenic and liver CFUs in wild type versus *IFNAR*^−/−^ mice (51). However, a more recent study has shown a higher burden of *Brucella* in wild type mice compared to *IFNAR*^−/−^ mice, indicating that Type I interferon response is detrimental to the host (6). Resistance to *B. abortus* in the *IFNAR*^−/−^ mice was accompanied by elevated production of IFN-γ and NO, and decreased apoptosis compared to wild-type mice. Although type I IFN served as a useful indicator for STING activity in this study, ultimately, the experience with the *IFNAR*^−/−^ mice suggest STING is controlling infection through Type I IFN-independent mechanisms.

*Brucella* potently inhibits apoptosis, contributing to chronic infection; however, the mechanisms behind this process are unknown (30, 52). By down-regulating STING and subsequent IFN-β production, *Brucella* could be actively inhibiting apoptosis that is dependent upon Type I IFN signaling. Further, by up-regulating miR-24 which in turn down-regulates Bim, *Brucella* could be avoiding UPR-mediated apoptosis, which is partially dependent upon Bim in other experimental systems (53). *B. melitensis* infection robustly induces CHOP, which is an upstream activator of Bim (20, 54). We did not detect a reliable effect of the anti-miR-24 on host cell apoptosis or cell death (data not shown). One likely explanation is that there are unidentified miR-24-independent mechanisms that inhibit apoptosis independently of STING and Bim down-regulation. Indeed, our previous RNAseq data (14) suggested that *Brucella* suppresses the expression of multiple pro-apoptotic molecules. Another possibility is that the other microRNA at the *mirn23a* locus counteract the miR-24 Bim mRNA targeting during infection (55).

In summary, this is the first report in which an intracellular bacterial pathogen evades full activation of innate immunity by down-regulating STING via miR-24 induction. It is noteworthy that a single miRNA species should have such a profound impact on a major cytosolic innate immune sensor and consequent *Brucella* replication. Our data may have implications for other important pathogens. For instance, miR-24 was up-regulated and cited as one of 7 significantly altered microRNAs controlling the transcriptional response to *M. tuberculosis* in macrophages (56). In a separate report, in transcriptomic data, *Tmem173* was suppressed by more than 50% at 4h and >75% decreased 12h following *M. tuberculosis* infection (57). Widely considered a “stealth” pathogen, *Brucella* can evade immune surveillance and persist chronically in macrophages (58). In contrast to this idea of *Brucella* as “flying under the radar”, previous reports have described *Brucella* subversion of toll-like receptor signaling via Btp1/TcpB (59-61). In this study, a single miRNA species exerted a profound effect on a major cytosolic innate immune sensor, with striking impact on intracellular replication. The data presented here present a critical mechanism by which *Brucella* actively sabotages cytosolic surveillance by the innate immune sensor STING to establish its intracellular niche.

## Acknowledgements

We would like to thank Dr. John-Demian Sauer for providing us with the *v-raf/v-myc* immortalized murine bone marrow-derived macrophages and Glen Barber for providing the STING^−/−^ mice. Bruce Klein supplied MyD88−/− femurs and Richard Dahl the mir23a −/− mouse bones. Conceptualization, M.K.,J.A.S., J.S.H., G.A.S., S.C.O., G.B., Methodology, M.K., J.S.H., R.D., J.A.S., Investigation, M.K., J.S.H., Y.L., E.G., J.W.T., D.G., T.H., F.C., Statistical analysis, J.E., Writing—Original Draft, M.K. Writing—Review & Editing, M.K., J.S.H., S.C.O., G.A.S., J.A.S., R.D. Funding Acquisition, M.K., S.C.O., G.A.S., J.A.S., G.B.

## STAR Methods

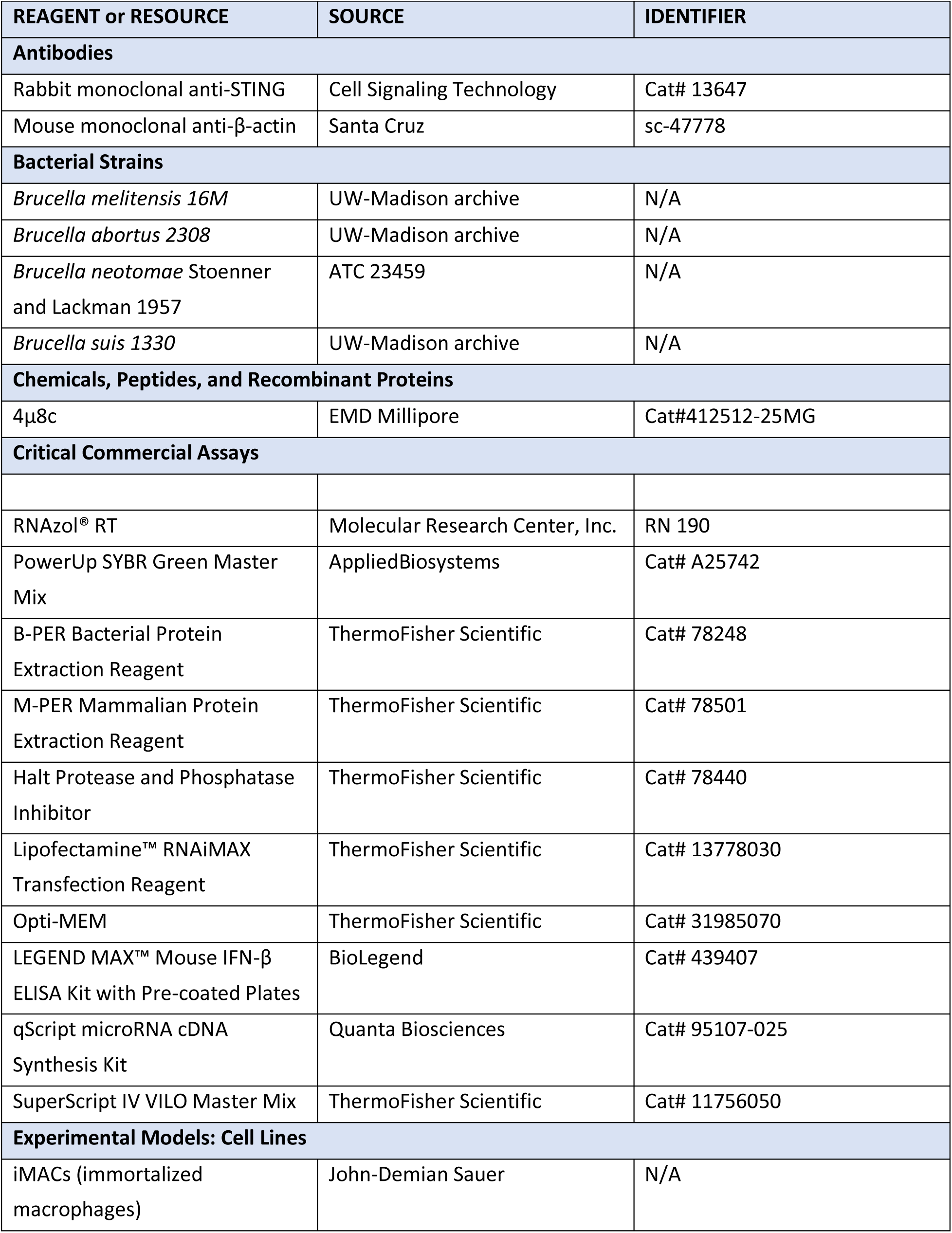
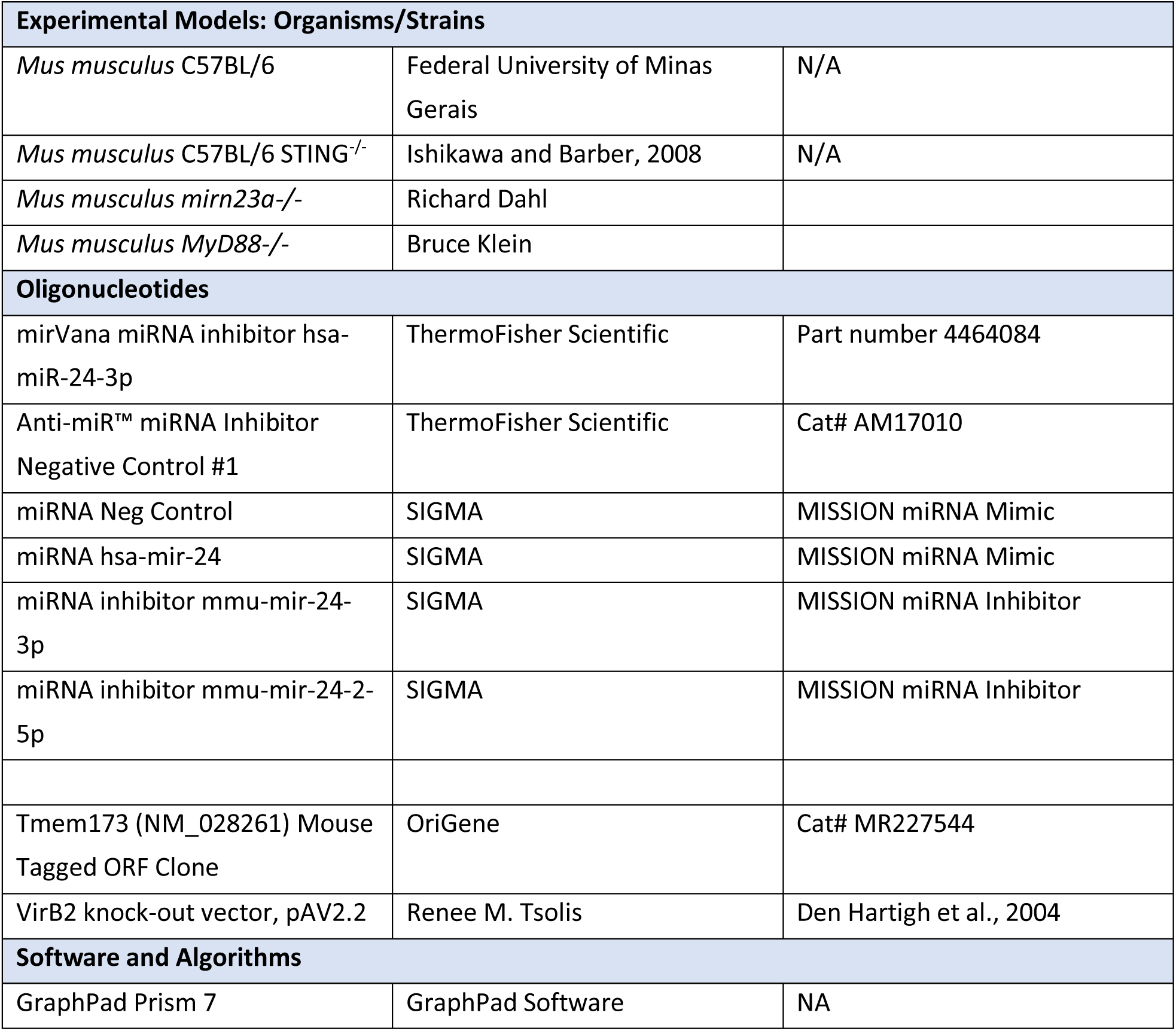

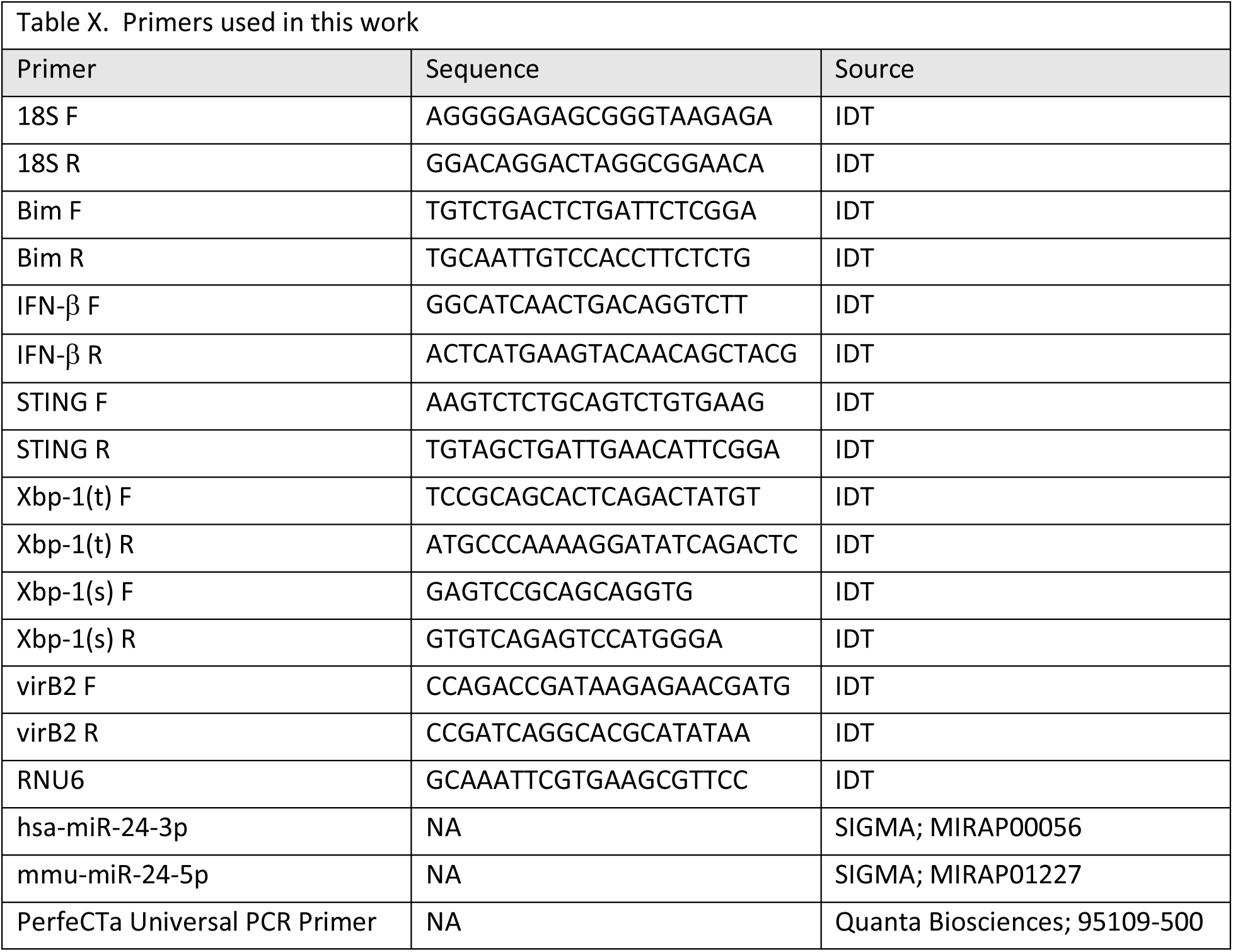

## Contact for Reagent and Resource Sharing

Further information and requests for resources and reagents should be directed to by the corresponding author, Judith A. Smith

## Experimental Model and Subject Details

### Mice and Ethics Statement

Wild type C57BL/6 mice were purchased from (resource?) and housed at the Federal University of Minas Gerais (UFMG). STING (*Tmem173*)^−/−^, mice were described previously (10). All mice were maintained at UFMG and used at six weeks of age. All animal experiments were pre-approved by the Institutional Animal Care and Use Committee of UFMG (CETEA #128/2014).

*In vivo* infections in mice were carried via intraperitoneal injection of 10^6^ CFU of *B. abortus* virulent strain S2308. Five female C57BL/6 and 4-5 STING^−/−^ mice were infected with *B. abortus* and sacrificed at three or six weeks post-infection. To count *Brucella* CFU, individual spleens were macerated in 10 ml saline, serially diluted, and plated in duplicate on *Brucella* Broth agar. After 3 days of incubation at 37°C, the number of CFU was determined as described previously (30).

### Mammalian Cell Lines

*V-raf/v-myc* immortalized murine bone marrow derived macrophages (BMDM) were a generous gift from Dr. John-Demian Sauer at the University of Wisconsin-Madison. These macrophages were from C57BL/6 mice. *V-raf/V-myc* immortalized BMDM were generated in our lab from leg bones of *MyD88*^−/−^, STING^−/−^, and *mirn23a*^−/−^ mice obtained from researchers listed in the table above. These mice all had a C57BL/6 background. All immortalized macrophage cell lines were cultured at 37°C with 5% CO_2_ in RPMI supplemented with 1mM Na pyruvate, 0.05mM 2-mercaptoethanol, and 10% FBS. Apart from Figure 1, immortalized macrophages were used for *in vitro* experiments.

### *Brucella* strains

*Brucella* strains *B. melitensis, B. abortus, B. suis*, and *B. neotomae* were from archived stock of the University of Wisconsin-Madison and cultured using Brain Heart Infusion broth or agar (Difco) at 37°C. All experiments with *Brucella* strains were performed in a Biosafety Level 3 facility in compliance with the CDC Division of Select Agents and Toxins regulations according to standard operating procedures approved by the University of Wisconsin-Madison Institutional Biosafety Committee.

VirB2 deletion mutants of *Brucella* were derived through homologous recombination following the methods described by den Hartigh et. al., 2004 using the plasmid pAV2.2 (generous gift from R.M. Tsolis). Briefly, exponentially growing *Brucella* were made electrocompetent following standard microbiological methods. Electrocompetent *Brucella* were then electroporated with pAV2.2 and VirB2 deletion mutants were selected for kanamycin resistance and carbenicillin sensitivity. VirB2 deletion was confirmed in these clones using PCR.

In experiments using heat-killed (HK) *Brucella* as a control, inactivation of *Brucella* was as follows. After growing *Brucella* in culture, the sample was quantitated using spectrophotometry. *Brucella* was then aliquoted in microcentrifuge tubes and placed in a 56°C waterbath for 1 h.

### Infection Assays

Immortalized macrophage cell lines were plated on 6-well tissue culture plates at 0.4 × 10^6^ cells per well in 2 ml culture media. The next day, the media was replaced, and cells were infected with 100 MOI *Brucella* determined by spectrophotometry (OD at 600 nm) through a formula established by a *Brucella* growth curve. Cells were incubated at 37°C. Four hours post-infection, cells were washed 3x with 2 ml/well warm PBS and fresh media containing 10 μg/ml gentamycin was added. Incubation continued until time points indicated for the experimental assays. In some experiments, the IRE1 endonuclease inhibitor 10 μM 4μ8c was added to the cultures 1 hour prior to infection.

### Colony-Forming Unit (CFU) assays

Cells were washed 3x in PBS to remove extracellular bacteria. Then, 1 ml of cell lysis buffer (dH_2_O + 0.1% Triton X-100) was added per well. CFU were determined by serial dilution plating on BHI agar after 3–4 days.

### Quantitative Polymerase Chain Reaction (qPCR)

Total cellular RNA was processed using RNAzol RT reagent (Molecular Research Center, Inc.) following the manufacturer’s protocol. Then cDNA was prepared from either mRNA, using the Superscript IV VILO system (Invitrogen) or microRNA using the qScript system (Quanta biosciences). Samples for Quantitative PCR were analyzed using SYBR Green and the delta-delta Ct method to calculate relative fold gene expression using a StepOnePlus thermocycler (ABI). Endogenous genes used for comparative expression were either 18S (mRNA) or RNU6 (microRNA). The primers used in this study were designed using IDT’s online primer design tool or purchased.

### Quantification of IFN-β by ELISA

Culture supernatants from cells were collected and frozen at −80°C until assayed. A mouse IFN-β ELISA kit was used following the manufacturer’s protocol. Absorbance at 450 nm and 570 nm were determined utilizing a BioTek microplate reader. Quantitation of mouse IFN-β was determined by standard curve.

### MicroRNA Inhibitor and Mimic Transfection

Macrophage cell lines were seeded on 6-well tissue culture plates at 0.4 × 10^6^ cells/well. The next day, miRNA mimics (miR-24-3p), miRNA control, and miRNA inhibitors for miR-24-3p were diluted to 0.28 μM using Opti-MEM and cells were transfected using using RNAiMAX Reagent following the manufacturer’s protocol. One day after transfection, cells were infected as described above.

### Western Blot Assays

Cell lines (infected or control) were washed with PBS and scraped off the well then transferred to a microcentrofuge tube and pelleted (4k RPM, 5 min). Supernatant was completely removed and cells lysed with M-PER reagent (ThermoFisher Scientific) according to the manufacturer’s protocol. Whole-cell lysates were resolved by 12% SDS-PAGE. Samples were then transferred to polyvinyldene difluoride (PVD) membrane and immunoblotted with anti-STING primary antibody (ThermoFisher Scientific) and anti-β-actin primary antibody (Santa Cruz), followed by a fluorescence-conjugated secondary antibody (LI-COR). Proteins were visualized and quantitated with the Odyssey system (LI-COR).

### Quantification and Statistical Analysis

CFU values were and standardized mRNA expression levels were summarized in terms of means ± standard deviations and displayed in graphical format using bar charts, stratified by experimental conditions. Comparisons between experimental groups were conducted using two-sample t-test or analysis of variance (ANOVA). Pairwise comparisons between multiple groups were conducted using Tukey’s Honestly Significance Difference (HSD) method. Residual and normal probability plots were examined to verify the model assumptions. Linear regression and Pearson’s correlation analyses were conducted to evaluate bivariate associations. Statistical significance is indicated in the figures (* p<0.05, ** p<0.01, *** p<0.005, **** p<0.001, ns not significant). Statistical analyses were conducted using SAS (SAS Institute Inc., Cary NC), version 9.4.

