## Supplemental Figure 1 for "*Brucella* suppress innate immunity by down-regulating STING expression in macrophages"

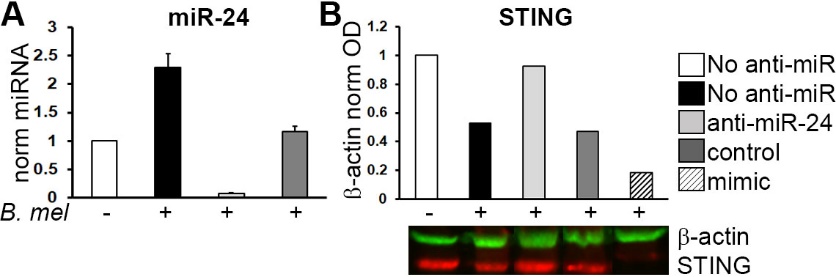

**Figure S1. miR-24 and STING expression in anti-miR and mimic transfected cells**. A) macrophages were not transfected or transfected with anti-mi24 inhibitor (I) or control miR (Ctrl). Cells were subsequently uninfected (NI) or infected with 100 MOI *B. melitensis*. After 24h, cells were harvested for RNA and miR-24 levels determined by qPCR with normalization to RNU6 and uninfected control. Note, miR-24 levels in the infected negative control transfectants were not as high as the non-transfected infected controls, but significantly higher than in specific anti-miR-24 treated cells. B) Macrophages were transfected with control microRNA, anti-miR-24-3p or miR-24-3p mimic. 24h following infection with *Brucella*, cells were lysed and whole cell lysates resolved by SDS-PAGE. STING and β-actin were detected using western blot and immunofluorescence. Blot is representative of 2 experiments.
